# Using Stakeholder Insights to Enhance Engagement in PhD Professional Development

**DOI:** 10.1101/2021.04.28.440824

**Authors:** Deepti Ramadoss, Amanda F. Bolgioni, Rebekah L. Layton, Janet Alder, Natalie Lundsteen, C. Abigail Stayart, Jodi B. Yellin, Conrad L. Smart, Susi S. Varvayanis

## Abstract

There is increasing awareness of the need for predoctoral and postdoctoral professional development and career guidance, however many academic institutions are only beginning to build out these functional roles. As a graduate career educator, accessing the vast silos and resources at a university and with industrial partners can be daunting, yet collaborative endeavors and network development both on and off campus are crucial to the success of any career and professional development office. To better inform and direct the efforts of graduate career offices, forty-five stakeholders external and internal to academic institutions were identified and interviewed to gather and categorize perspectives on topics critical to career and professional development offices. Using a stakeholder network visualization tool developed by the authors, stakeholder engagement can be rapidly assessed to ascertain areas where offices have strong connections and other areas where additional efforts could be directed to enhance engagement. General themes from interviews with internal and external stakeholders are discussed to provide graduate career educators with various stakeholder subgroup perspectives to help prepare for successful interactions. Benefits include increased engagement and opportunities to collaborate, as well as the opportunity to build or expand graduate career development offices.

## Introduction

Institutions of higher education hold myriad potential connections between predoctoral and postdoctoral researchers, faculty and administrators, internal university offices, industry partners, professional societies, and funding organizations. Internal university partnerships are vital, ranging from predoctoral and postdoctoral researchers themselves and university faculty, to externally-facing communications and alumni development offices. University career and professional development (CPD) programs also develop and rely on external partnerships, particularly with programming and resources designed for predoctoral and postdoctoral researchers. While CPD programs understand that these partnerships improve the predoctoral and postdoctoral training experience, provide pipelines for entry of predoctoral and postdoctoral researchers into the workforce, and lead to synergies and collaboration, the full value of these relationships may not be completely understood to internal and external partners. Our work explores the foundational value of internal and external intersections and how to best leverage them to prepare predoctoral and postdoctoral researchers for the workforce. The aim is to more efficiently and successfully coordinate relationships that meet all stakeholders’ needs with a more thorough understanding of stakeholder objectives and the relative value of engagements.

This project is a spinoff of the National Institutes of Health Broadening Experiences in Scientific Training – NIH BEST (National Institutes of Health & Office of Strategic Coordination, 2014) Consortium’s Annual Meeting (2018). The Consortium (active 2014 – 2019) was comprised of programs at 17 higher education institutions challenged by the NIH to develop innovative approaches to prepare predoctoral and postdoctoral researchers for a wide range of careers in the biomedical research enterprise. The Consortium’s final Annual Conference (2018) explicitly invited collaborations, presentations, and conversations through joint programming with institutions beyond the Consortium – ranging from well-established pioneer predoctoral and postdoctoral professional development program institutions to newer and aspiring institutional programs interested in establishing professional development programs, as well as external private and non-profit collaborators. This research project emerged from the massively audacious goal of the group that “all predoctoral and postdoctoral scholars have support and resources needed to explore and pursue all careers, and faculty and institutional leadership buy-in to the importance of this mission.”

The goals of this publication are to bring awareness within the higher education community about various stakeholders that commonly engage with graduate CPD, shed light on stakeholder perceptions of career development and engagement, broaden the composition of engaged collaborations, and provide engagement tools. This information can help institutions and individuals quickly self-assess and visualize strengths/opportunities with stakeholders for the purposes of CPD at their institution, and over the long-term build stronger relationships and partnerships.

### Defining stakeholders

The authors quickly realized that to attain their goal, a broad set of stakeholders would need to be consulted. Informed by the authors’ experience in industry relations and engaging with higher education, stakeholders were identified, classified, prioritized, and consolidated into a rapid tool for stakeholder engagement (see methods). Table 1 displays internal and external stakeholder classification groups and subgroups: internal stakeholders include pre- and postdoctoral researchers, faculty/administrators, and external-facing staff; external stakeholders include non-profit and society partners and industry employers. Each stakeholder subgroup was approached with a specific set of questions to explore their perceptions of graduate career education and professional development, as well as their motivations for engaging with any of the other stakeholder groups, ultimately seeking advice for how to best engage them in CPD programming.

**Table 1:** Stakeholder classification and examples

| Stakeholder Group | Sub-Group Interviewed | N = 45 | Examples |
| --- | --- | --- | --- |
| Internal Stakeholders | 1) Pre- and postdoctoral Researchers | 9 | Predoctoral students, Postdoctoral researchers |
|  | 2) Faculty/Admin | 8 | Assistant, Associate, Full Professors, Chairs of Department, Directors of Research Centers |
|  | 3) External-Facing Staff | 12 | Staff administrators with roles in career services, industry / government relations, technology transfer / licensing, communications |
| External Stakeholders | 4) Non-profit/society Partners | 8 | Trade organizations, professional societies, non-profits, business associations |
|  | 5) Industry Employers | 8 | Small and large companies, intellectual property firms, consultancies, accelerators |

Descriptions of desirable and required skills, as well as resources to share with pre- and postdoctoral researchers were sought out.

### Internal Stakeholders

The group that frequently receives the most focus from CPD professionals includes pre- and postdoctoral researchers. Fostering stakeholder engagement from pre- and postdoctoral researchers is critical for providing effective programming (Sherrer & Prelip, 2019). CPD programs less frequently look beyond pre- and postdoctoral researchers to engage a full spectrum of university stakeholders within their own ecosystem.

Foremost among them is the faculty. Data from the BEST Consortium indicated that a large majority of faculty are supportive of career training for various careers and have recognized that pre- and postdoctoral researchers participating in CPD activities were happier and making timely progress toward degree completion – a fact that can be used to recruit additional internal stakeholder engagement from faculty (Brandt et al., 2020; Chatterjee et al., 2019; Watts et al., 2019). Faculty don’t always believe they have the knowledge or resources to assist pre- and postdoctoral researchers whose career interests lie outside academia, although they largely support their career pursuits (Engelhardt & Alder, 2020; St. Clair et al., 2019; Varvayanis et al., 2020; Watts et al., 2019). Promoting transparency, encouraging the normalized need for career support, and recommending conversations to initiate CPD coaching will help bridge some knowledge and awareness gaps.

Identifying internal collaborators with external facing roles has the potential to reduce or obviate the need to develop *de novo* program components, to seed ideas that leverage each other’s networks and knowledge domains, and to overcome common roadblocks (Meyers et al., 2016). Cost-sharing on resources and events are an added benefit for engaging with internal partners (including student leaders), and opens doors for pre- and postdoctoral researchers to engage in on-campus job shadowing and internship opportunities for experiential learning (Meyers et al., 2016; Sherrer & Prelip, 2019; Van Wart et al., 2020). Collaborations with alumni relations and development offices can dramatically expand the network of potential speakers and mentors (Qualls et al., 2021); it can also lead to coordinated fundraising campaigns for CPD initiatives. Moreover, establishing relations with other internal partners with external-facing roles in industry or federal relations, technology transfer, licensing, or research commercialization can amplify the skill sets of pre- and postdoctoral researchers seeking experiential learning opportunities in real-world settings (Van Wart et al., 2020).

### External stakeholders

Efforts of career development professionals must simultaneously be internally and externally focused to fully understand the skills current pre-and postdoctoral researchers need to execute an informed transition into careers of their choice (Sinche et al., 2017). In addition, external focus identifies potential employers to build pipelines for these researchers, attract funding sources, access training opportunities to support their CPD, and increases visibility and accessibility of the resources offered by CPD programs. External stakeholders include partners and employers as seen in Table 1. The categorizations in Table 1 are based on how CPD practitioners primarily interact with each group but can overlap with other categories, as many partners are also employers.

Stakeholders in external groups such as industry, non-profits and government agencies, including professional societies and associations, have long partnered with academia in disseminating research and technical training. They increasingly offer skill-building and career development opportunities via conferences and webinars to assist current pre- and postdoctoral researchers in the career selection process. Additionally, societies acknowledge multiple career options and wield positional influence to support culture change within academia.

A primary function of CPD programs is to strengthen the future workforce by preparing pre- and postdoctoral researchers for interaction with external stakeholders, ultimately, future employers. Therefore, the foci of these outward-facing efforts should be strategic to broaden networks and facilitate connections. Engaging external employer stakeholders in networking events, site visits, job shadowing, internships and panel discussions makes it possible for pre- and postdoctoral researchers to explore and test-drive various PhD careers (Fuhrmann et al., 2011; Meyers et al., 2016; Van Wart et al., 2020). This study provides useful tools and insights to address stakeholder engagement.

## Methodology

### Stakeholder Identification & Engagement

#### Study Design

This study is intended as an exploration of a field, and an opportunity to observe emerging phenomena. The research team identified both internal and external stakeholder groups relevant to graduate CPD programs in order to further identify values that each stakeholder places upon bidirectional interactions to advance CPD for pre- and postdoctoral researchers. Common themes were identified through open-ended questions and unanticipated value propositions to develop potential approaches for improving interactions and methods of engagement between the university and potential partner organizations.

#### Data Collection

Interviews offer a rich and robust capture of perspectives (Lofland & Lofland, 1995), providing in depth, open-ended responses and opportunity for dialogue between researcher and participant. Single-interaction, semi-structured interviews, conducted either in-person (before COVID-19), by phone, or online via Zoom, were used with standardized questions and optional probing follow-up questions as needed (see Appendix A for complete interview question list). Once core questions were established, subsets of parallel but slightly amended questions relevant to each stakeholder group were developed. Due to privacy concerns, neither recording nor transcriptions were approved for human subjects. Instead, interviewers took digital or manual live notes. In all interviews, verbal consent was sought from participants.

#### Recruiting/Access to and selection of participants

Selection criteria for interview subjects were designed to be representative of identified stakeholder groups, with an initial goal of including 5-6 individual interview participants per stakeholder subgroup. External stakeholders initially included industry and non-profit/societies grouped together, but after the first few interviews, the research team discussed the difference in themes that arose from these interviews and arrived at the consensus to classify them as two distinct groups. Therefore, additional participants were recruited to ensure there were 5-6 participants for each of these subgroups.

Potential participants were identified by several means: by each interviewer independently, based on individuals known to the interviewers; referrals within and across the authors’ networks; or vetted Google searches. The invitation selection process considered a broad variety of types of organizations or groups based on stakeholder classification (Table 1), and identified a sampling of individuals within a particular subgroup that included a variety in perceived support for the premise, academic disciplinary background, education level, as well as across social identity categories including gender, race/ethnicity, and international status. Other factors considered included leadership/experience level, tenure in respective organizations, and any experience or interest in working at the interface of professional/career development for varying purposes. This background was not known for all and some were purposefully naïve or underexposed to CPD initiatives in higher education. A review of prospective participants’ general characteristics served to uncover similarities or duplications. Efforts were made to ensure that a variety of types of organizations were represented in each subgroup.

#### Conducting interviews

Once selected, prospective participants were invited by email to participate in a short 15-20-minute interview. The template invitation (Appendix B) included a brief script of the study’s purpose and description, plus an overview of the questions to be asked. Interviewer and participant found a mutually convenient time and format (in-person, phone, or video conference call).

After identifying stakeholder groups and subgroups (see Table 1 and Discussion for the importance of flexibility and refinement), four interviewers conducted a total of 45 stakeholder interviews (of 55 invitations). Themes were collected separately for the groups that used the same sets of questions to differentiate responses (i.e. pre- and postdoctoral researchers versus faculty and administrators, external partners versus employers). Following the first round of interviews, one group had a higher sample size than the others, therefore, to keep group numbers roughly equal, target recruitment goals were updated to a minimum of eight interviews per subgroup (Supplemental Table 2).

#### Data analysis and interpretation/validity

A multi-stage process was used for data analysis and interpretation, including sorting of sensitizing concepts (Blumer, 1954), and analysis and reduction of data through application of grounded theory (Glaser, B. G. & Strauss, 1967; Strauss & Corbin, 1994), leading to the identification of emergent themes, hierarchical grouping, and concept categorization. One member of the research team who did not conduct any interviews was designated as coder. The coder and each individual interviewer reviewed and ‘binned’ potential initial themes emerging from keywords and phrases, and separated text into categories using paraphrased concepts or the original words from each participant into each row of a spreadsheet with category column headings. This ongoing collaborative synthesis of data and collection of emergent themes contributed to the iterative data reduction and display process, including a process of contrast/comparison, and noting patterns and themes (Miles & Huberman, 1994).

At the conclusion of coding of each interview, coder and interviewer reviewed the initial data-sorting and ‘binning’ to ensure themes were appropriate and consistent with participant intent. With each subsequent interview within a stakeholder group (internal, external) and subgroup, themes were refined; new ‘bins’ were created if the participant’s comments did not fit into an existing bin. If a response fit into two themes, then they were placed in both bins and coded as repeated. A second text review after all interviews were complete was conducted by the coder, with all authors working collaboratively in a process of inter-rater reliability. The team appraised the interviews in the larger context to make sure the original interview notes and emergent themes were not in conflict, still represented participant viewpoints, and to catch any themes missed in the first review. A final complete review by all authors prior to summation repeated this process through robust group discussion and collaborative decision-making (Kaner et al., 2014). All final themes and comments were reviewed and consolidated to assure researcher agreement on the accuracy of the themes and statements were selected to represent each theme.

Unique themes found in interview text were highlighted and reported as representative themes that arose when multiple instances of each theme occurred within a stakeholder subgroup. Themes with fewer than five mentions by participants are not incorporated in the Results text, and are available in Appendix C; however, themes with three or four mentions are included in the figures.

#### Subjectivity/Ethical Issues/Limitations

All interviews were conducted by researchers who are professionals in higher education (e.g., program directors, associate directors, assistant deans) strongly invested in CPD for pre- and postdoctoral researchers (within offices of graduate education, postdoctoral affairs, CPD programs, evaluation). All interviewers are professionally full-time employed women in the US, and the study team included US and international interviewers, both people of color and white.

The possibility of selectivity bias exists, in opinions or stories shared by participants based on their roles, and in the selection of participants within individuals’ networks tending toward supporters of graduate CPD. To attain a balanced view of various stakeholder subgroup perspectives as well as in recruiting participants, three methods were used to avoid compounding selectivity bias: 1) a semi-structured interview style with pre-selected questions (see Recruiting). 2) explicit recruitment of “nay-sayers” as well as “supporters” of university/organizational partnerships and graduate career training. 3) Google searches to identify further participants beyond known networks, as well as requests to members of known networks to suggest individuals the authors had no previous connection to, who might not be interested in or knowledgeable specifically about graduate professional development, but were in positions related to industry-university engagement activities. Each interviewer conducted interviews with individuals they knew and those they had never met, from offices or organizations they were familiar with and those with which they had no prior knowledge. Specific inclusion of the question, “Do you follow the national conversation about career development and outcomes of PhD-trained scientists?” helped determine their level of awareness of the subject matter. Nonetheless, the authors recognize the need to interpret findings with caution and that they will not represent all possible viewpoints. The results should be viewed as pilot data to inform additional research.

All participant names are anonymized using pseudonyms and all data is de-identified prior to sharing in accordance with IRB approved protocols (Rutgers – FWA00003913 Study ID Pro2020222400; PittPRO STUDY19110306-I4; UNC IRB# - 19-3054; Boston University H-40210). Note that the pseudonyms were randomly assigned and their perceived ethnicity or gender is not intended to represent the individual participant. Any correlation with a theme and gender, race or ethnicity is purely coincidental. Demographics were gathered at the last step to fill in post-analysis to prevent unconscious bias or revealing of identity or demographics of any participant. In the results, only pseudonyms are used, using the code in Supplemental Table 2.

#### Stakeholder Engagement Tool

The authors observed a gap between the perceived awareness of the variety of stakeholders and the ability to assess and capitalize on strengths of potential existing partnerships with internal and external stakeholders. To help rapidly assess the two, a tool was created. During the development of the stakeholder engagement tool, categories were refined based on discussion among the authors, their combined experiences working with various stakeholders, as well as a two-way influence of the interview process (the tool influencing the interviews, and the stakeholder perspectives from the interviews influencing refining the tool).

The authors first shared the tool publicly at an international conference hosted by the Graduate Career Consortium. Input from conference attendees participating in that workshop helped question previous assumptions and helped to better describe how users would customize the tool.

## Results

A total of 45 individuals were interviewed by four interviewers (Supplemental Table 1). Consenting participants consisted of both men (42.2%) and women (57.8%). Participant demographics were US (71.1%) and international (28.9 %), African American (15.6%), Asian (17.8%), White (62.2%), and Hispanic (4.4%). Participants were geographically diverse across the US, with interviewers accessing their own networks primarily across the Northeast, Southeast, and Midwest United States, but also extending geographic representation via national organization contacts and referrals.

Stakeholder organizations included public, private, and public-private hybrid institutions of higher education (some with medical schools); large and small companies in or serving the biotech, medtech or pharma industries; as well as foundations, non-profit organizations or business associations serving STEM fields. Summaries of the major themes arising from each set of stakeholders along with specific examples from representative responses are presented in decreasing order of frequency of mentions across the interviews.

### Stakeholder 1 – Internal Predoctoral Students and Postdoctoral Researchers

These interviews (Appendix A) probed attitudes towards devoting time to CPD. Responses from ‘frequent users’, ‘occasional users’, or ‘non-users’ of CPD programming were separated.

#### Support for and perceived benefits of CPD programs identified by pre- and postdoctoral researchers

##### Theme 1.1: Efficiency, productivity, and content (12 mentions)

Users find CPD affects productivity in a positive way and allows for pre- and postdoctoral researchers to get a broad overview of resources. Glenn elaborated on the need to focus on tools that pre- and postdoctoral researchers can use and apply to their lives currently. Occasional-user Gretel expressed the desire for more pre- and postdoctoral researchers’ input in the design of CPD activities. Non-users, such as Gunnar, thought that these events were a waste of time spent on common sense information that could be obtained from their lab more efficiently.

##### Theme 1.2: Networking, community and role models (11 mentions)

Community-building advantages of professional development activities, e.g. bridging pre- and postdoctoral researchers across labs, were discussed by researchers such as Gaston and Gael. They also noted the benefit of activities that allowed for networking with hiring managers and recruiters rather than only networking with other scientists who do not have influence in non-academic spaces. This theme aligns with findings of a sense of community with pre- and postdoctoral researchers attending professional development training, especially in cohort or mandatory participation models (Sherrer & Prelip, 2019). Some users also mentioned the confidence-building, cultural, and gender-based inclusivity that these programs can provide via intentional conversations and making role models prominent.

##### Theme 1.3: Exposure and decision-making of career paths (6 mentions)

Some pre- and postdoctoral researchers felt that they needed to decide if a particular career was for them or not. International users, such as Guangli, expressed a need for this programming, citing lack of familiarity with careers in the US. A non-user, Gunnar, who plans an academic career, acknowledged that it is important to understand industry priorities and needs to help them guide their own future pre- and postdoctoral researchers who seek industry positions.

##### Theme 1.4: Embedded/Required (6 mentions)

Requiring professional development activities in the curriculum would allow for consistent messaging each year of training. Gaston commented that if it is required, then pre- and postdoctoral researchers will need to make time for the activities. Both frequent users and non-users thought embedded programming will provide support and motivation in their career exploration, and this is echoed in a report generated by a separate multi-stakeholder workshop (Bixenmann et al., 2020). One of the non-users, Gunnar, pointed to business schools that successfully integrate CPD in curricula. Indeed, many business schools have good examples of externship/internships required for curriculum (Van Doren & Corrigan, 2008).

Other perceived benefits include: Prestige for recruitment (4 mentions) [see Appendix C for details]

#### Opportunities to improve identified by predoctoral and postdoctoral researchers

##### Theme 1.5: Consistent exposure throughout training (18 mentions)

Pre- and postdoctoral researchers communicated the desire for consistent exposure to CPD activities throughout training. Gaston, a frequent user as a postdoctoral researcher, mentioned having “rueful regrets” in retrospect about not participating in professional development during PhD training. Many commented that the best approach is regular interactions starting early in training, for even a few hours a month.

##### Theme 1.6: Centralized access (5 mentions)

Predoctoral and postdoctoral researchers clearly voiced their desire for centralized programming, as they believe it to be an important and positive feature. Representative across several users, Gael captured this sentiment, noting that centralized CPD programs equalizes access to resources and institutionalizes the concept of career development to facilitate acceptance by faculty. Gael also raised a concern regarding equitable access in the absence of centralized programming; for instance, faculty advisors may not be knowledgeable enough to help direct pre- and postdoctoral researchers interested in careers outside of their own path, or select graduate programs may create separate professional development activities accessed by only their predoctoral and postdoctoral researchers. Hence professional development programming at the school or university level is more desirable.

Other challenges with fewer mentions include: topics around growth/challenges (4 mentions), and the express need for faculty permission (4 mentions) [Appendix C]

**Figure 1.**
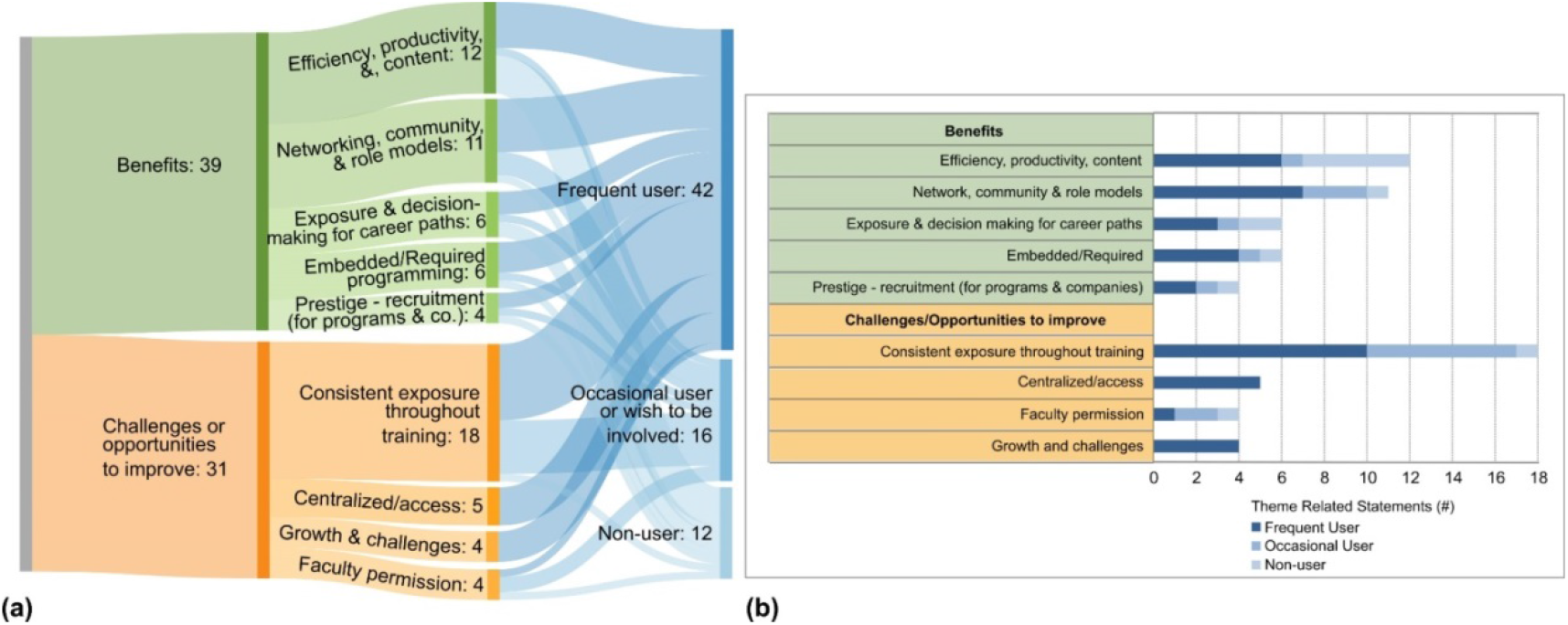
Themes from Internal Pre-and Postdoctoral Researchers (a) Sankey diagram and (b) stacked bar graph representing the same data of the number of mentions for each theme representing benefits and challenges/opportunities to improve mentioned by frequent users, occasional users and non-users.

### Stakeholder 2–Internal Faculty and Administrators

Participants answered the same questions as the Internal pre- and postdoctoral researchers subgroup above (Appendix A) focused on probing attitudes towards devoting time to CPD, and opinions of such opportunities (existing, or hypothetical) available to pre- and postdoctoral researchers. Themes that arose naturally divided participant responses into categories that were later identified as ‘enthusiastic supporter’, ‘cautious supporter’, and ‘non-supporter’ responses.

#### Perceived ‘benefits’ of CPD identified by Faculty/Administrators

##### Theme 2.1: Evolving training requirements and climate (12 mentions)

Faculty feel, as Fariba mentioned, that they are living vicariously through their pre- and postdoctoral researchers by being excited for those who are successful in a variety of career paths. The benefit ranged from acknowledgement that training grant applications require a description of career development activities to how participation in these activities can improve mental health. Faculty and administrators acknowledged institutional peer pressure nationally among top tier universities to provide these opportunities, and that it is the right thing to do for the pre- and postdoctoral researchers.

##### Theme 2.2: Awareness of workforce outcomes (9 mentions)

Among enthusiastic supporters, there was an acknowledgement that there are limited faculty positions available and that pre- and postdoctoral researchers are choosing a variety of career paths. Frank commented that it is valuable to expose researchers to all one can do with a PhD and that it is helpful to be transparent with workforce outcomes of alumni as examples for current pre- and postdoctoral researchers. However, less supportive respondents believe that professional development is not related to graduate education, that it de-emphasizes academia as a career path, and that individuals who leave academia, cannot return.

##### Theme 2.3: Promotes career exploration and planning (6 mentions)

Faculty and administrative stakeholders noted the value of equipping predoctoral and postdoctoral researchers with a breadth of training beyond the academic career path, and that providing role models in different jobs allows current researchers to see themselves in those positions in the future. Fariba said PhD alumni, who are successful professionals, have mastered skill sets in their jobs, and hence hearing from them can add value to training and career exploration. These stakeholders also noted how mandatory Individual Development Plans (IDPs) set the framework for helping pre- and postdoctoral researchers organize their goals to obtain the skills necessary for those jobs.

Other benefits include: Cycle of positive fulfilment for program (3 mentions).

#### Opportunities to improve identified among faculty and administrators

##### Theme 2.4: Tailored experience and exposure (17 mentions)

Multiple comments were made about the need for students to be exposed to options early in graduate school. Fariba believes that graduate programs should encourage researchers to think about career planning before they start their graduate careers. It was also suggested that time commitment and type of engagement should align with educational stage. Others thought that for professional development activities to be efficient, it should be modular so the training experience can be tailored to each pre- and postdoctoral researcher’s interests and can be prioritized individually.

##### Theme 2.5: Concerns and perceptions (13 mentions)

Some faculty saw professional development as a waste of time that could lengthen time to degree completion. Finley’s comment indicates the belief that pre- and postdoctoral researchers need to do one thing really well, even if only to focus on their dissertation research. Finley also believes that professional development, although important, should be done outside of research training, as researchers do need to do a lot of soul searching. Fariba and other faculty showed concern that smaller universities would not be able to launch their own professional development programs.

##### Theme 2.6: Narrow definition of professional development (6 mentions)

Several of the more cautious and non-supportive participants interviewed thought that professional development only revolved around academic skills. Finley, for example, believes that research development is career development. Furthermore, Finley believes that as educators, faculty and administrators should teach students to recognize their limits, and that their training is a privilege. These individuals did not indicate awareness that the phrase ‘professional development’ includes more than learning to publish, teach, and write grants.

**Figure 2:**
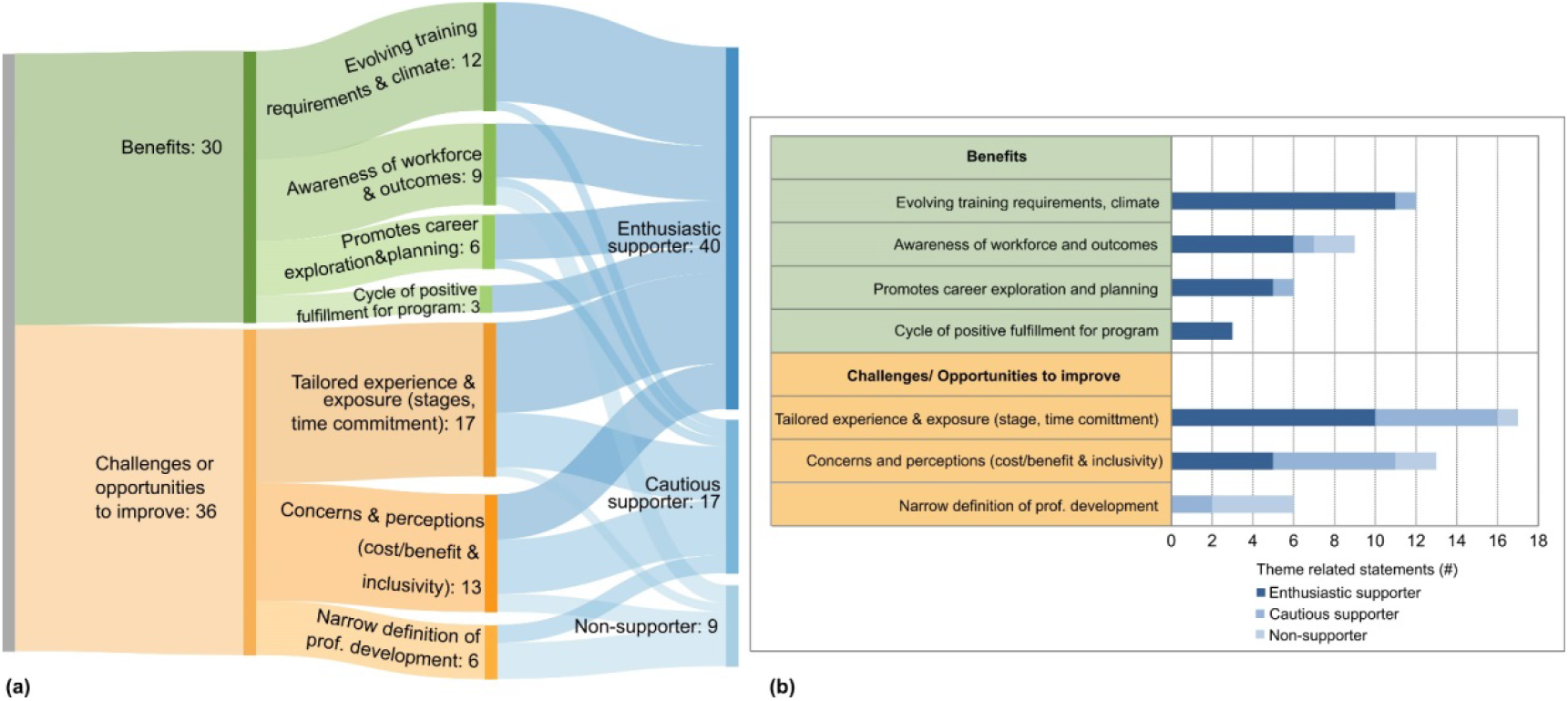
Themes from Internal Faculty/Administrators (a) Sankey diagram and (b) stacked bar graph representing the same data of the number of mentions for each theme representing benefits and challenges/opportunities to improve mentioned by frequent users, occasional users and non-users.

### Stakeholder 3: External Facing Staff (Industry Relations, Tech Transfer, Communications, Alumni Relations and Development)

Interviews with these stakeholders were guided by a different set of questions (Appendix A). The goals of these questions were to identify with whom external facing offices at institutions typically interacted, and around what topic(s) the majority of their interactions centered. While these offices are open to all institution affiliates, do many of their external contacts have an interest in STEM pre- and postdoctoral researchers?

Respondents in external facing offices interact with small and large businesses and foundations (8 individuals), investors (4 individuals), alumni (3 individuals), inventors (2 individuals), and innovation centers (2 individuals). Views from several other affiliations or identites include experts, educators, presenters, business development and intellectual property professionals, an institution’s business school (e.g. on consulting projects), grateful patients, hospital systems, government, and professional organizations. It is of note that the themes below do not include a federal relations point of view.

#### Reasons to engage with industry identified by external-facing offices

##### Theme 3.1: Innovation and entrepreneurship (16 mentions)

The interaction between industry and academia toward innovation and entrepreneurship is captured by Simha’s comment that she needs to bring the voice of the market to the university, helping to inform the direction of academic research to provide added value. The growth of biomedical technologies has spurred the enthusiasm to work with scientists to translate research and technologies. These offices work closely with researchers at the institutions, including pre- and postdoctoral researchers, as well as with industry representatives to evaluate the viability of projects.

##### Theme 3.2: Building partnerships (11 mentions)

Examples of partnership benefits include fundraising, increasing awareness of internship programs, and involving industry in courses or events developed for pre- and postdoctoral researchers.

Interactions with these groups leads to a better understanding of how small and large companies can better support initiatives such as providing hand-on experiences to these researchers. Shandra explained that these partnerships matter, as large companies can guide content creation of curriculum as well as provide longer-term help to universities through the development of ideas and resources. Soren encouraged the more holistic approach, and coordinating with state-wide and economic development offices to create partnerships between academia and industry.

##### Theme 3.3: Fundraising or financial support (10 mentions)

Shyla shared that another primary purpose for external-facing offices to interact with campuses is to create sponsored research. Interestingly, Shanice emphasized mutually beneficial goals leading to fundraising, where their external-facing office’s motto is “time, talent, treasure”.

Another theme with fewer mentions included the perceived benefit to pre- and postdoctoral researchers (4 mentions) [Appendix C]

#### Reasons external stakeholders engage with academia identified by external-facing staff

##### Theme 3.4: Early access to emerging technologies and innovations (18 mentions)

A recurring theme centered on external stakeholders’ keen interest in early access to emerging technologies or innovation. Simha commented on the interest, describing external stakeholders’ intellectual curiosity, love of science, and being part of new discovery, while Saachi’s opinion shared external stakeholders’ interest in getting first access to emerging technologies developed in academia. Sree advocated for using industry partners’ academic interests to incorporate experiential learning experiences within curricula.

##### Theme 3.5: Developing an entrepreneurial mindset (11 mentions)

Another recurring theme was the goal of assisting in developing an entrepreneurial mindset, with Shandra’s response capturing why industry partners are interested in PhD level researchers – because people who are adaptable, flexible, and can work in teams are needed across the workforce. They recommended instilling an entrepreneurial mindset to make pre- and postdoctoral researchers more competitive. They also advocated that these skills can and should be taught to all students to better serve both themselves and the workforce more broadly, referring to a joint research brief on entrepreneurial mindset (Ernst & Young & Network for Teaching Entrepreneurship (NFTE, 2018), and to reports published by the EU Commission (Bacigalupo et al., 2016) and World Economic Forum (Drexler et al., 2014)

##### Theme 3.6: Faculty expertise or connections (8 mentions)

External offices perceive faculty expertise or connectors as a strong motivator for external partners to interact with universities. Shandra indicated that companies can get advice on how to improve their businesses and smaller companies can connect with support services.

##### Theme 3.7: Talent identification (6 mentions)

Unsurprisingly, external facing offices believe some external stakeholders view campuses as a talent identification hot-spot to build a workforce pipeline. Sahana points out that employers are seeking talent but are not interested in career fairs, implying that they would rather vet talent by getting to know the pre- and postdoctoral researchers in more organic ways than career fairs allow.

##### Theme 3.8: Provide scholarships/grants, or fund programs (5 mentions)

External- facing staff believe that external partners are interested in funding research at a variety of levels, including generating revenue from patents or licensed products. Sven emphasized sponsored research programs as a way for companies to provide funding for universities through support of faculty projects.

Another less mentioned theme includes: Pay it forward (3 mentions).

#### External stakeholders’ interest in STEM predoctoral and postdoctoral researchers identified by external-facing staff

##### Theme 3.9: Business experience and STEM knowledge (10 mentions)

Among external- facing staff, there was a strong sentiment that business experience was more valuable than technical background. Shandra noted entrepreneurial mindset skills are important preparation for any career. Although STEM knowledge is important, the distinguishing feature for some companies is business acumen.

##### Theme 3.10: Expertise/talent pipeline (10 mentions)

According to the external facing staff, one major benefit for external companies to interact with their office is access to new talent. Saachi indicated that a side effect of technology application review is a connection with predoctoral researchers who are potential talent. The predoctoral and postdoctoral researchers who provide the expertise on collaborations and consultation are also possible future employees at these companies.

##### Theme 3.11: Mutual Benefit (10 mentions)

The idea that predoctoral and postdoctoral researchers’ relationship with other internal and external stakeholders is mutually beneficial was reflected by multiple stakeholders. Sahana noted that their institution’s technology transfer internship program helps students get jobs, publicizes the university, and helps fulfil external companies’ employment goals.

Other themes included: Support student needs (4 mentions).

#### External-facing staff’s additional thoughts

##### Theme 3.12: Specific advice to predoctoral and postdoctoral researchers (11 mentions)

Many had suggestions for researchers to better prepare for their careers, but important advice was summed up best by Scott, who said that everyone should be prepared to learn on the job because the value of a PhD is that one can learn something new independently. Taking risks, trusting your critical thinking, and relying upon other transferrable skills developed in training, were strong messages shared by the external facing staff.

##### Theme 3.13: New perspectives on training needed (6 mentions)

These internal stakeholders believe that to best prepare for any future career, it is helpful for predoctoral and postdoctoral researchers to expose themselves to many different experiences. Scott thinks it is important to gain skill sets to prepare for future interactions, no matter what career plan one has. Thus, embedding industry perspectives on workforce development into the curriculum can facilitate job seeking as well as establish long-term interactions between industry and academia.

Other themes with fewer mentions included: the need for a communication coordinator (4 mentions), and an alumni engagement role (3 mentions) [Appendix C].

**Figure 3.**
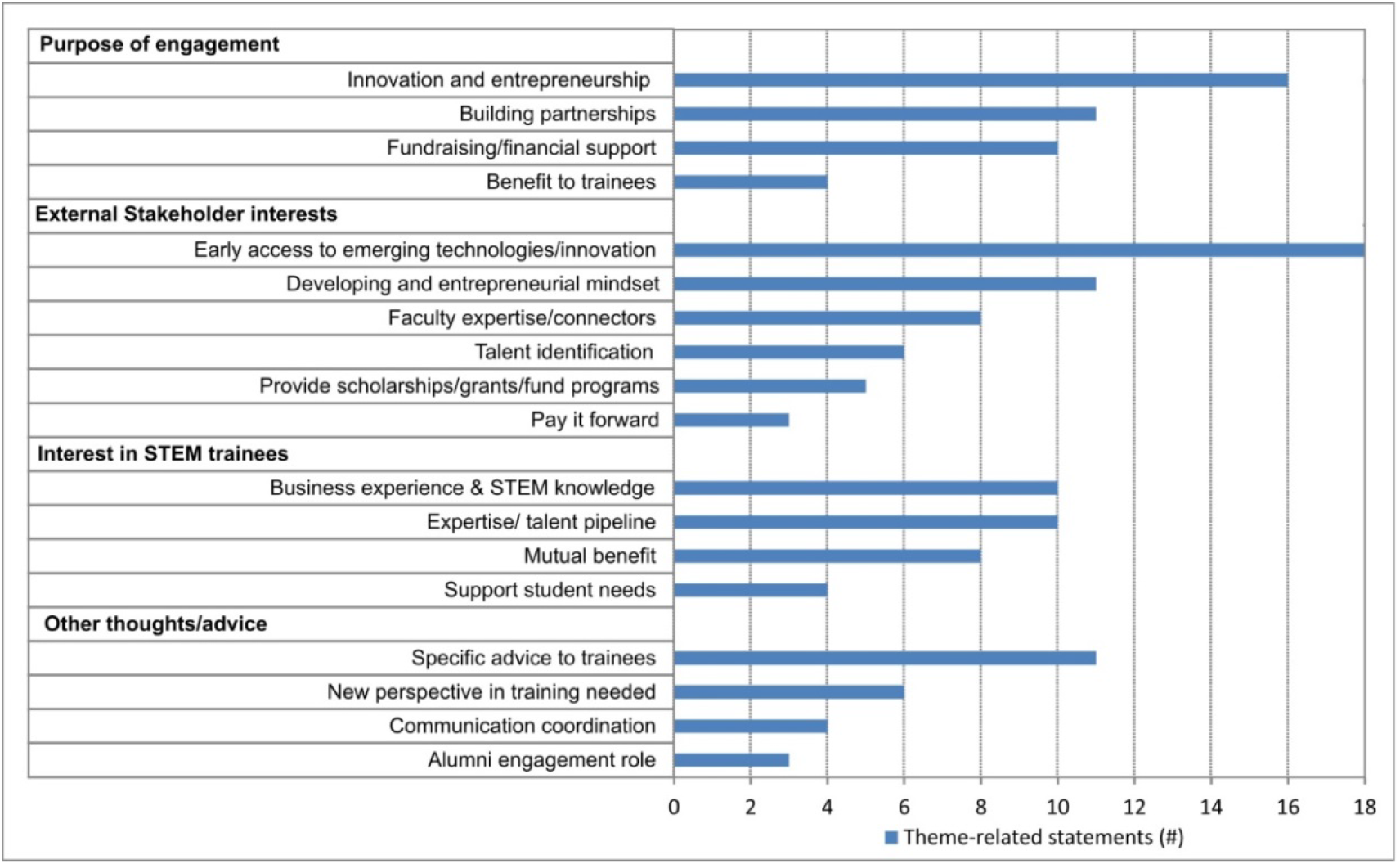
Themes from External-Facing Staff: Bar graph representing the number of mentions of each theme.

### Stakeholder 4: External Partners – Societies, Foundations, Non-Profits

Interviews with societies, foundations, and non-profits indicated that there were many partnerships and exchange of services by which they interact with academia including: networking and community building, providing resources for honing career skills, generating scholarship and publications, facilitating advocacy, providing feedback and advice, and creating funding opportunities.

Discussion with the stakeholders from societies, foundations and non-profits brought new themes to light. These highlight many available resources that are not always accessed by institutions. The most common reason societies, foundations, and non-profits provided for wanting to engage with academia was to build relationships to connect academics with the mission of their organization.

#### External partner organizations’ engagement with academia

##### Theme 4.1: Building relationships (16 mentions)

Building relationships was a large motivator for societies, foundations and non-profits to interact with academia. Ellen sees the role of the non-profits as the access point bringing academia and the private sector together. Forming bridges and connections is viewed as essential for the non-profit’s missions.

##### Theme 4.2: Prestige, recognition, public visibility (8 mentions

While these external partners want to visit and interact with academia, they see it as a status-raising opportunity. Ellen says they are interested in the impact it can have on their organization, such as receiving public credit or being included as a full partner in endeavours. Non-profits are interested in benefits of prestige and visibility their organization could receive for such engagement.

##### Theme 4.3: Catalyst for connections, knowledge (5 mentions

Foundations and societies see themselves as a space for understanding commonalities and allowing for synergistic relationships to form. Eric captured this sentiment, commenting that their goal is to create a flow of knowledge between universities. This theme indicates that societies, non-profits, and funders may have more resources and opportunities available that are underutilized by universities.

#### Societies, foundations, and non-profits resources to offer

##### Theme 4.4: Expertise and advice (6 mentions

Societies and foundations have several scientists on staff possessing a wide range of expertise and perspectives. Ellen expressed that scientific staff should be tapped for their expertise, while Evan indicated they can provide the context for scientists to be understood by governmental agencies, as well as provide scientists context on how policies are made.

Additional resources with fewer mentions included online resources, guides (4 mentions), experiential learning (4 mentions), and self-exploration (2 mentions) [Appendix C].

Challenges faced by external partner organizations were few: funding (4 mentions), connecting with target audience (3 mentions), and flexible, creative models and the need for culture change (3 mentions) [Appendix C].

#### External partner organizations’ view of career preparedness improvements

##### Theme 4.5: Develop and broaden skills, learning approach (11 mentions)

The need for pre- and postdoctoral researchers to expand their skills was discussed frequently by external partners. Evan specified the importance of diversifying training to include non-science courses and to expand researchers’ skills to include science communication and persuasive writing.

##### Theme 4.6: Take early initiative (8 mentions)

Many external partners stressed the need for pre- and postdoctoral researchers to take initiative early to create opportunities and explore their options. Ellen voiced the advice succinctly when she advocated for researchers to get involved in an organization and develop relationships well in advance of their job search.

##### Theme 4.7: Improve self-efficacy and growth mindset (6 mentions)

Societies, foundations and non-profits encourage pre- and postdoctoral researchers to adopt a growth mindset, and to be aware of demand for their skills. Ebony impressed upon not constraining oneself, and to explore and grow with one’s science and research. Others touched upon the necessity to acknowledge how opportunities offered by societies are equally important and valuable to research internships.

Other themes with fewer mentions include: Listen with humanity, broaden diversity (4 mentions), networking and engagement opportunities (2 mentions), get involved in professional societies (1 mention), and defining success (1 mention) [Appendix C].

#### External partners’ view of engaging with academic institutions

##### Theme 4.8: Invitations (5 mentions)

These non-profit and society stakeholders are keen to engage and to disseminate their resources. As Ellis highlighted, these organizations are accessible, and they encourage universities to invite them to visit campus and coordinate their visits. Their view is that universities should reach out and contact the non-profits rather than solely relying on the reverse.

Another theme included inviting researchers off campus (2 mentions). Again there were notably fewer themes of challenges identified by this stakeholder subgroup, all with fewer mentions, such as offerings should be integrated into training (2 mentions), to engage at all levels with academic institutions (2 mentions), that the wrong people are making decisions (1 mention), and the need for more industry mentors.

**Figure 4.**
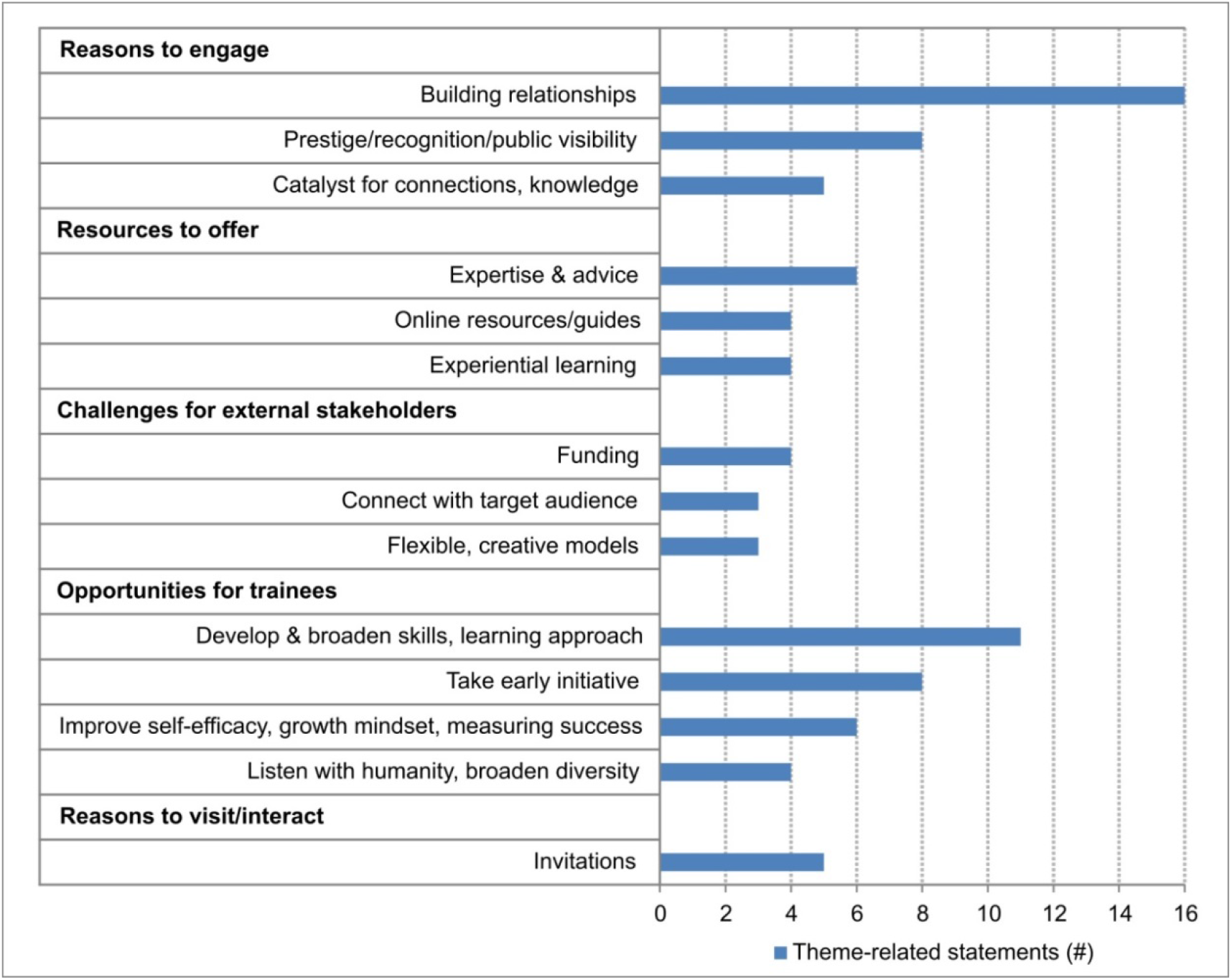
External Partners: Societies, Foundations, Non-profits: Bar graph representing the number of mentions of each theme.

### Stakeholder 5 – External Employers – Small and large companies, intellectual property firms, consultancies, accelerators

External employers interviewed for this study include large pharmaceutical, biotech, government or national labs, consulting firms, intellectual property firms, policy or communication organizations as well as small business/startups, accelerators and boutique consulting agencies.

Types of engagement already in place with universities include: recruiting events, career fairs, tours/site visits, case studies/workshops, serving on advisory boards for curriculum development, and internships. A key goal for this kind of engagement is to maintain relationships.

A representative sampling of interviews with external employer stakeholders revealed additional themes and underscored themes already brought to light in the previous interviews.

#### Reasons why industry external stakeholders engage with academia

##### Theme 5.1: Recruiting and broadening their reach (8 mentions)

Companies are interested in being involved in professional development programs. As Iris explained, these programs are a pipeline into the company, even if the company is small and only a few PhDs are hired. Some companies see a relationship with a university as a geographically local recruitment tool.

##### Theme 5.2: Building long-term relationships (5 mentions)

Shandra shared that the same person from the company went to an institution every year, thereby creating a strong connection. She believes that this helps develop a sense of trust and community, allowing these representatives to then serve as technical recruiters to help pre- and postdoctoral researchers prepare for interviews. When an alumna/us acts as the liaison, there is an added benefit of helping maintain connections and creating a sense of community.

Other themes with fewer mentions included: the need for bidirectional partnerships (4 mentions), and advisory and feedback roles (3 mentions). [Appendix C]

#### Training and collaboration industry-academia partnership benefits

##### Theme 5.3: Alumni and mentoring (8 mentions)

The modes of interaction between industry and academia include alumni working with their *alma mater*. Of note, Ivan indicated that the personal recruiting via doctoral alumni is not used enough. Other interactions, such as pre- and postdoctoral researcher outreach, were touched upon. For example, Ivan provided feedback that their company appreciates it when these researchers initiated relationships.

Importantly, if a company’s needs are already met, they admitted to having no motivation to interact with academia, as they believe that their reputation alone was sufficient to draw job applicants.

##### Theme 5.4: Programs and industry expertise (6 mentions)

Irina pointed out that industry has efficient resources to attack bigger problems, where academia has complementary skills for broad questions. Ian reiterated the need for increased awareness of industry operations, such as drug development and marketing.

Other themes include: Learn industry-relevant skills (4 mentions), Recalibrate importance (3 mentions), and Grants and academic collaborations (3 mentions).

#### Differing priorities and organizational complexities – challenges with industry

##### Theme 5.5: Differing values (5 mentions)

Ira believes that academia should welcome collaborations and not distrust industry standards as industry is highly regulated. Furthermore, Ira advised that academics should lose their negative attitude toward non-technical roles. Scientists who do go down this path should be equally celebrated as successful. The different value systems between industry and academia should not create a level of disdain between the two.

Another theme with few mentions includes reference to a desired single point of contact (3 mentions).

#### External industry employers’ views on challenges with predoctoral and postdoctoral researcher preparedness

##### Theme 5.6: Understanding options, industry culture and priorities (8 mentions)

Industry professionals’ advice to pre- and postdoctoral researchers is, as Ivan shared, the importance of understanding what an industry job entails and what the job title means. Imani stated that observation or hands-on experience is critical before applying.

##### Theme 5.7: Develop communication skills (8 mentions)

A frequently mentioned theme highlighted the need for pre- and postdoctoral researchers to develop good communication skills before embarking on a career in industry. Imani stressed upon the need to better synthesize information and communicate complicated topics in an exciting and concise way.

##### Theme 5.8: Present experience and motivation (7 mentions)

While it is well-acknowledged that PhD-trained individuals have good training and experience, it is important to be able to translate this appropriately. Irina advised that these researchers need to understand the problem-solving process is valuable, and that they should not panic about what they don’t know in the industry setting.

##### Theme 5.9: Relationship building and collaborations (5 mentions)

Collaborations are seen as essential, and hence the ability to build and sustain relationships is critical to succeeding in industry. Ivy emphasized the need to help PhDs understand the collaborative nature of industry research. Additionally, Ira was quick to point out that each industry employee is of equal value: marketing managers and scientists are equally important team members.

Other themes with fewer mentions included: Faculty culture change (4 mentions) [Appendix C].

#### How to encourage industry professionals to interact and visit campus

##### Theme 5.10: Invitations (6 mentions)

Several companies are glad to visit or interact with academic institutions. Ivy highlighted that they do this when students invite them for career panels, informational interviews, campus visits, or to set up site visits. Simply being asked to be on a career panel or other programs is sufficient.

Other themes included: High-impact events (3 mentions), Match-ups (3 mentions), and Open-minded to industry (3 mentions), preference for hosting site visit at company (1 mention), and challenge of limited time and resources to engage with academia (1 mention).

The vast majority of large and small external employers interviewed were unaware of the national conversation about career development and outcomes of PhD trained scientists.

**Figure 5.**
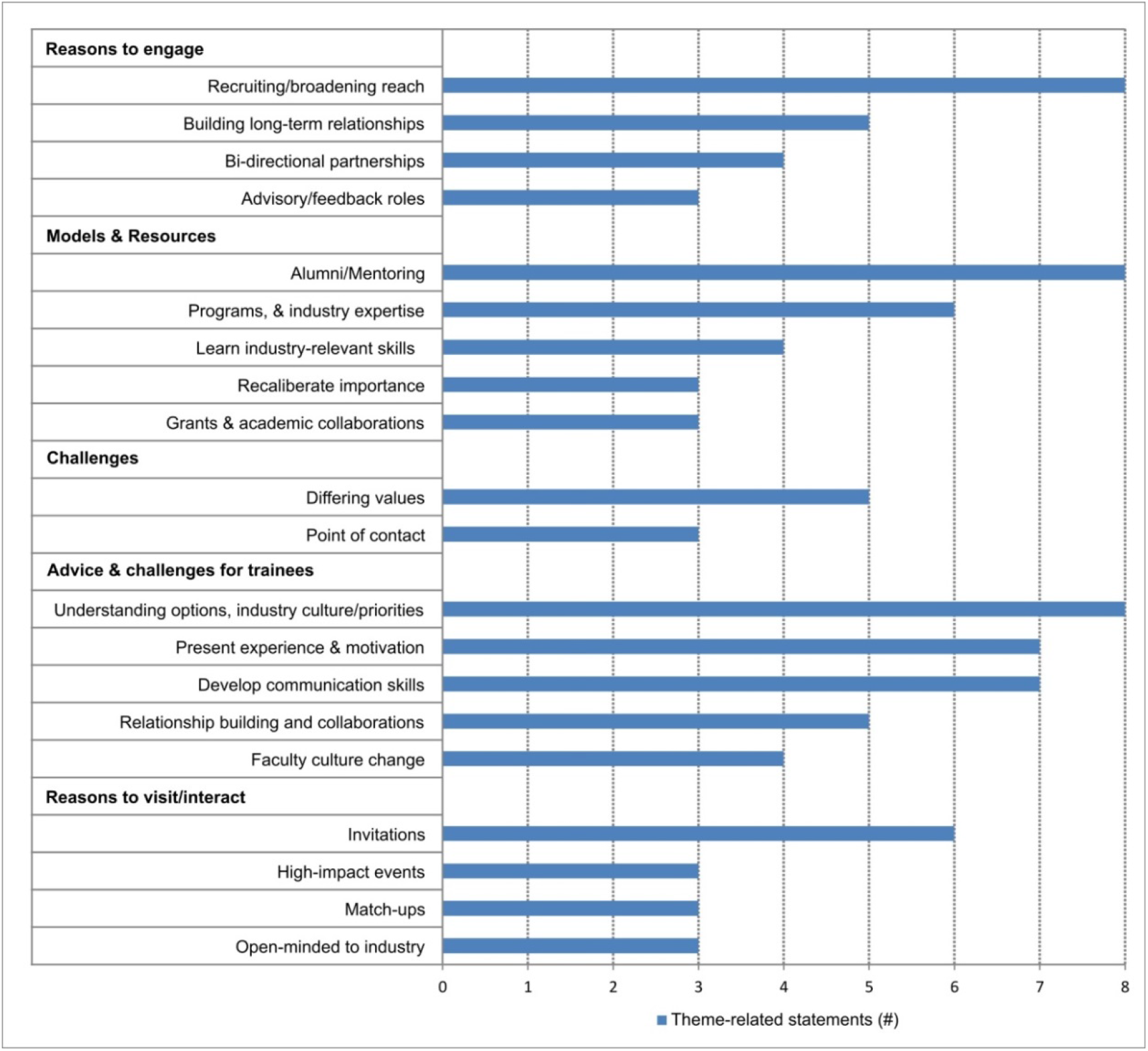
Themes from External Employer Stakeholders – Industry – Large and Small Businesses: Bar graph representing the number of mentions of each theme.

#### Stakeholder outreach tool purpose, use and actions to take

A stakeholder engagement tool was created for readers to quickly identify which stakeholder group they are primed to interact with most efficiently. The rapid assessment tool can help practitioners quickly focus on existing strengths at their institution on which they can rely, as well as on areas of improvement and possible links to approach stakeholders strategically. Coupled with the themes found in the interview data, a targeted approach can be developed to improve stakeholder engagement.

The rapid assessment tool is fully customizable to reflect local organizations with whom to partner and those that already might have existing relationships. A 360 degree view of perceived stakeholder engagement can be quickly determined by encouraging colleagues around campus to fill out the tool. The resulting scores will inform discussion across offices to see where perceptions align and where there might be differences in scores. Since the tool is easy to use and automatically creates a visual output by summing scores in each quadrant of the ensuing graph (representing internal pre-and postdoctoral researchers, faculty/administrators; external-facing staff; external partners; and external employers) practitioners gain a quick, holistic view of their engagement.

There are three basic approaches to action as a result of filling out the stakeholder engagement tool as seen in Table 2 below. See the supplemental file to download and use the stakeholder engagement tool.

**Table 2:**
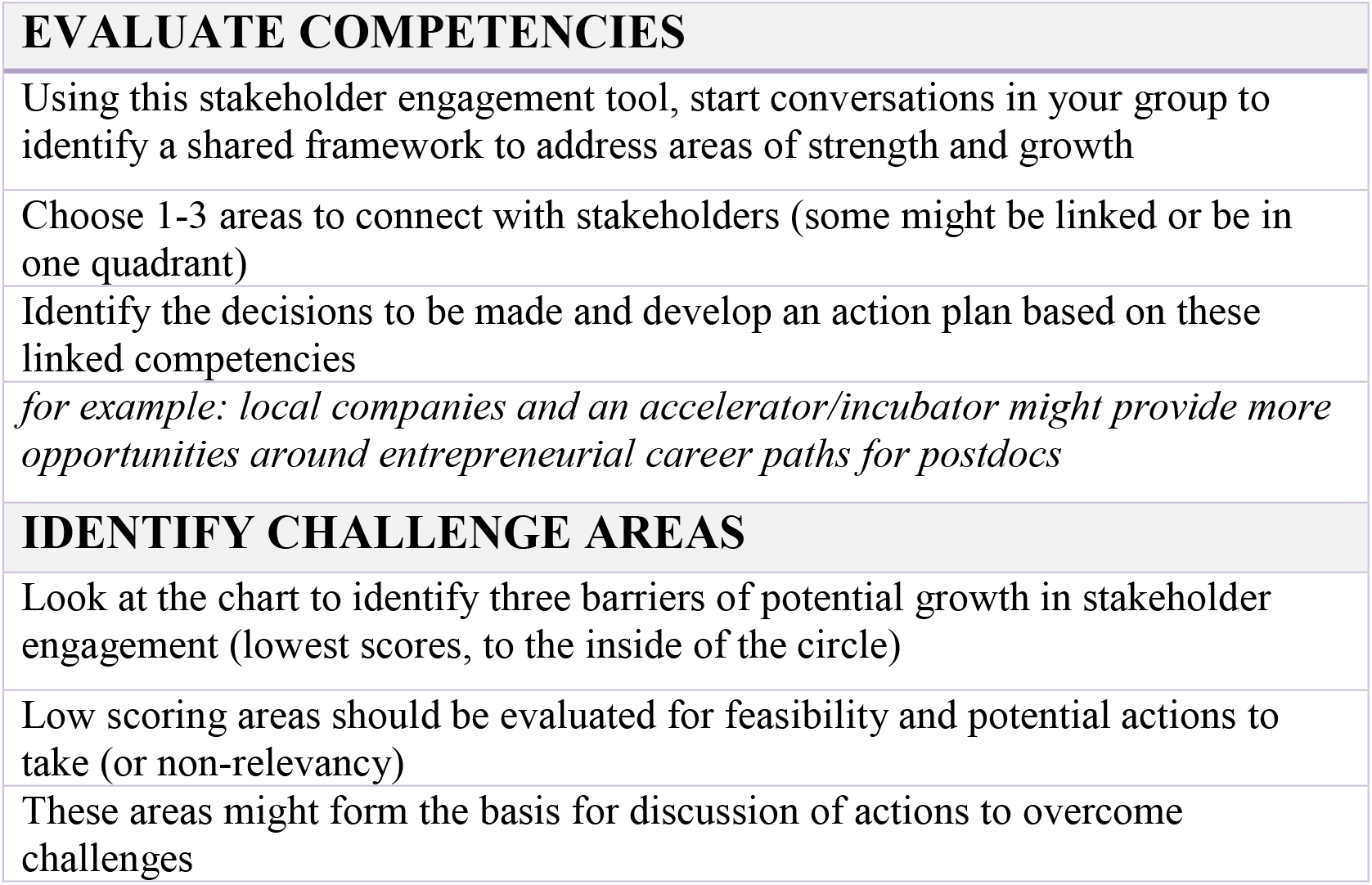
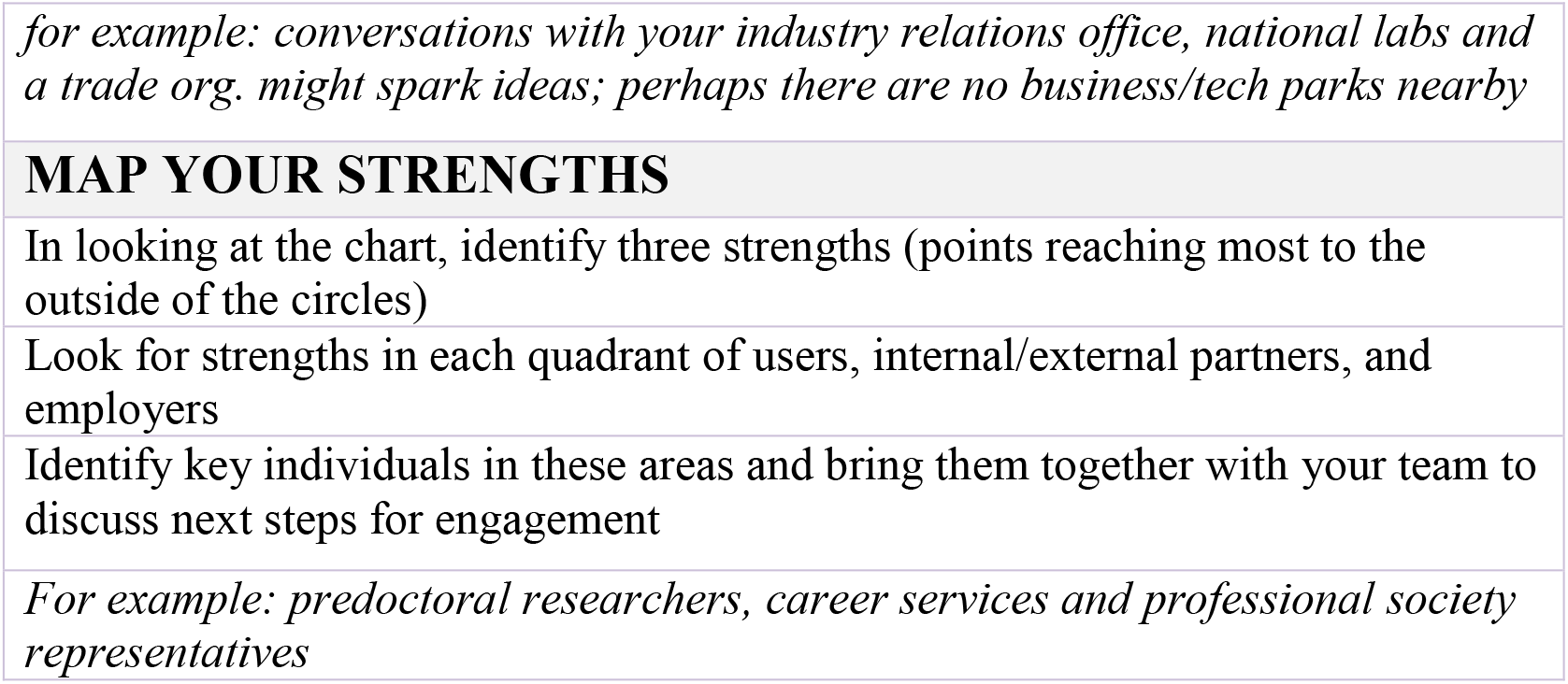
Three approaches to action following use of the stakeholder engagement tool

**Figure 6:**
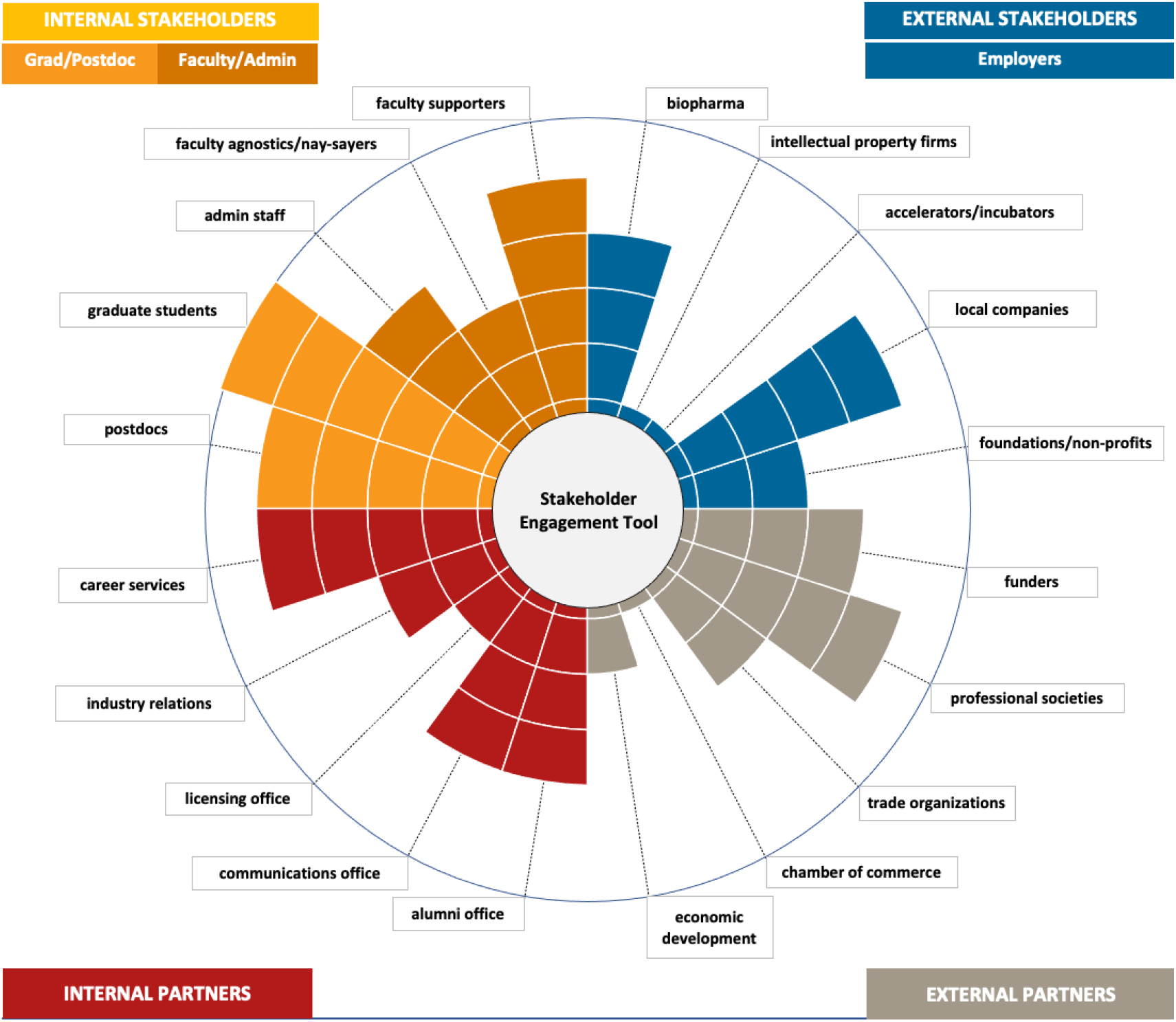
Stakeholder Engagement Tool: A rapid assessment tool for internal and external stakeholders to evaluate competencies and determine strengths for engagement in career and professional development programming [see supplemental file to download and use the tool]

## Discussion

### Common Themes

When considering the diverse stakeholder groups, common themes emerged across the different internal and external groups. Themes that reappeared across the various stakeholders suggest that these topics should be priorities for CPD practitioners when attempting to engage broadly.

There are individuals from all stakeholder groups interviewed who recognized the value of CPD activities for pre- and postdoctoral researchers, despite different perspectives on similar concepts, with individual participants varying in levels of enthusiasm and commitment. For instance, there is a distinct need for CPD programs across stakeholder groups, who engage for varying reasons. Among internal stakeholders, most pre- and postdoctoral researchers believe these activities benefit their career growth, many faculty and administrators believe the programs strengthen researcher career development and benefits their mental health, and all external-facing offices are keenly aware of both the value to pre- and postdoctoral researchers and how these activities generate interest among external stakeholders. External partners and employers desire that researchers are well-prepared when entering the workforce and encourage CPD programs to partner with them on program development. Notably, external partners develop many resources to aid CPD activities for pre- and postdoctoral researchers and encourage them to be involved and to use these resources.

Networking, making connections, partnering, and collaborating are seen as crucial aspects of CPD across all stakeholder groups. Pre- and postdoctoral researchers are interested in opportunities to build and grow their networks to help form their future careers, and faculty/administrators understand this to be critical to researcher development. Non-profits and foundations encourage non-prescriptive models for graduate learning, which could open the doors for more engagement across stakeholders, and industry partners offer assistance in developing programming and internship opportunities. Financial arrangements can be mutually beneficial: universities can benefit financially from industry interactions to support their research, and collaborative commercialization of research benefits industry. Additionally, external stakeholders are interested in connecting with academic institutions to have early access to emerging science and technology, develop partnerships to grow their own priorities, and build a talent-pipeline. External employers also showed keen interest in collaborating with faculty, and recommend that academics engage with industry partners on joint publications to help highlight these partnerships in the media (Butts & August, 2018), and facilitate culture change regarding opinions on industry collaborations.

The timing and content of CPD activities was another important focal point with multiple subgroups (pre- and postdoctoral researchers, external-facing staff, external partners and employers) suggesting the need for integrated, persistent, embedded, and flexible access to these activities. In particular, pre- and postdoctoral researchers suggest that there is a need for consistent exposure to CPD throughout training, including by some infrequent users who believe that this programming should be woven into the curriculum for maximal benefit, just as professional schools do. Historically, the perception was that pre- and postdoctoral researchers should focus on their careers after they complete their training, but this is not ideal as it delays the workforce pipeline (Meyers et al., 2016). In addition, some faculty stakeholders do not see the value of CPD activities and expressed some concern about the time their pre- and postdoctoral researchers dedicate to these activities. However, recent evidence-based research has shown that participation in internships, career development programming, K-12 outreach programs or IRACDA programs does *not* lead to increased time to degree or decreased productivity (Brandt et al., 2020; Gamse et al., 2010; Rybarczyk et al., 2011; Schnoes et al., 2018). Pre- and postdoctoral researchers, faculty-administrators and external partners all noted the value of flexible programming to encourage pre- and postdoctoral researcher engagement. All stakeholders pointed to the need to keep updated on both researcher needs and workforce needs when planning, designing, and executing CPD activities.

An important challenge identified by pre- and postdoctoral researchers, external-facing partners, and external stakeholders was the need to expand the purview of scientific training to include skill development for a variety of careers. Alongside this, these stakeholders comment on the challenge to normalize CPD activities in the larger context of training, and the need to help academic leaders (e.g. faculty and administrators) understand that academic career preparation is only a part of CPD, and that other types of skill development are necessary, as they are complementary and important to the success of their pre- and postdoctoral researchers. For example, skills such as having increased self-efficacy is extremely beneficial to researchers when approaching their CPD (Bandura, 1993; Sherrer & Prelip, 2019). Relatedly, requiring faculty approval to participate in professional development activities could be a barrier and sets the tone that professional development is outside of the normal expected activities of a pre- and postdoctoral researcher. Despite agencies (e.g. National Institutes of Health, 2014) clarifying that pre- and postdoctoral researchers’ skill development is critical, the concept of career exploration has not permeated to all faculty and administrators – it is evident that for successful CPD implementation, practitioners need to be active in outreach and engagement with internal faculty and administrators (Meyers et al., 2016; Watts et al., 2019). CPD offices should work to identify new strategies for conveying their services and value to get faculty/administrator buy-in, e.g. more evidence-based research to convey the program’s benefits. Knowing the culture of local faculty, and attitudes toward CPD can help strategically design programs that will yield the highest number of participants (Meyers et al., 2016).

Among other challenges discussed, was the identification and implementation of streamlined methods to access or connect with the right resources or people. While pre- and postdoctoral researchers struggle to know where to locate resources at their institutions, suggesting the need for a centralized institutional CPD hub, external-facing staff commented on not knowing the appropriate people within academic institutions with whom to connect their external contacts. External stakeholders recommend universities have a visible “one-stop shop”, to encourage external partners or employers to connect with them. Additionally, external stakeholders note that it is critical for a graduate career office to have strong engagement between past and present CPD practitioners, to ensure continuity of relationships with the various stakeholders.

The authors’ knowledge of internal stakeholders allowed them to engage with a spectrum of users of CPD services including frequent users, occasional users and non-users, as well as a range of faculty/administrators showing enthusiastic, cautious or no support for CPD activities. The variety of internal stakeholders interviewed resulted in valuable conversations to identify where CPD practitioners can improve. For example, pre- and postdoctoral respondent interviews cited requiring an increased awareness of their needs and purpose for engagement to better align existing CPD opportunities and guide new ones, while faculty/administrator interviews highlighted perceptions of CPD offices, identified concerns, collected suggestions on tailored experiences and exposures, and identified the need for clearly defined CPD. It appears valuable for CPD practitioners to have a clear understanding of internal stakeholders’ needs and concerns to create effective programming. Simultaneously, university staff who engage with external stakeholders share similar interests to CPD offices that include supporting and giving advice to pre- and postdoctoral researchers. Hence collaborations with these partners can provide valuable external stakeholder perspectives.

### Stakeholder Engagement

The stakeholder engagement tool, when shared at an international meeting, helped a variety of practitioners evaluate and plot to extend their networks in a targeted and structured way. Feedback from university-based users included surprise to learn the number of partners that could be leveraged by sometimes small, understaffed offices tasked with serving large populations of graduate students and postdocs. Newly identified partnerships across campus as well as in the local community were seen as options not previously considered. Benefits cited by meeting attendees included the rapid ability to identify a wide variety of stakeholders with whom to work and partner to increase opportunities for their pre- and postdoctoral scholars. An additional value was the quick analysis of potential barriers to growth in stakeholder engagement that could guide future discussions to overcome challenges. The tool was seen as useful also because it can be tailored to each institutional setting, e.g. whether industry partners are in the local area, or if the university is a large, decentralized behemoth. Templates for how to reach out to potential partners were also reported to be useful. Of course, the tool is intended only as a first step in engaging stakeholders. More research and time investment is needed to determine the exact person to reach out to at various organizations if no existing partnerships exist, but the process is intentionally step-wise so that over time more stakeholders can be involved in CPD to benefit all parties.

CPD practitioners should consider engaging with both alumni and future employers as key stakeholders. Alumni are one of the most accessible external stakeholders for graduate career development work. The personal experience of alumni in the workforce provides critical input for assessing skills required by employers and to inform curriculum changes (Meyers et al., 2016; Sherrer & Prelip, 2019; Sinche et al., 2017). Furthermore, alumni connections are valuable for establishing external partnerships. Individual institutions vary in their ability to cultivate and engage alumni, influenced greatly by the existence of an alumni relations office with active engagement strategies, for example, social media sites and accessible directories (Qualls et al., 2021; Van Wart et al., 2020).

The process of identifying external employer stakeholders as well as engagement opportunities should include strategic consideration of the location of the institution. For example, universities located in urban areas with a high concentration of biopharma companies might develop mechanisms to promote pre- and postdoctoral researchers’ biomedical expertise, valuable to local external stakeholders (Collins et al., 2020; Van Wart et al., 2020). Universities who are more isolated might organize a conference or trek to a more biotech-rich area (Butts & August, 2018; Van Wart et al., 2020).

## Conclusions

This study brings to light fundamental career and professional development (CPD) concepts that span the various internal and external stakeholder groups interviewed. Learning from these opinions is valuable, and can help form recommendations in the creation, design, and sustenance of effective CPD activities at individual institutions. This study also presents the stakeholder engagement visualization tool, which can be used for rapid self-analysis of practitioners’ networks to assess strong stakeholder relationships and areas where the practitioner can strengthen their network. Coupled with the various themes from interviews with 45 internal and external stakeholders across the country in various roles, graduate career practitioners can use the themes presented as discussion points to interact with their own stakeholders to prepare for potential meetings with their various stakeholders. Meaningful and targeted engagement with various stakeholders is key to create and sustain successful graduate CPD programs.

## Appendix A: Rationale and Sample Questions for Stakeholders

### Project Rationale

In order to provide resources to predoctoral and postdoctoral researchers, we first need to determine whether our current understanding of ‘the value of a PhD’ is accurate, from the point of view of stakeholder groups; perhaps we are missing important aspects/beliefs that have not been adequately appreciated or explored. We will approach stakeholders and pose a specific set of questions that explore their relationship and vision of interacting with academia and, specifically, pre- and postdoctoral researchers and how we can better support those interactions. What we learn from them will help us to better support and develop relevant resources for our pre- and postdoctoral researchers.

### Stakeholder Groups and Questions

#### Internal stakeholders (Stakeholders 1 and 2)

a. Pre- and postdoctoral researchers
b. Faculty
c. Academic administration

##### Sequence of conversation and questions

i. Introduction: reminder of purpose
ii. Do you believe that career & professional development for predoctoral and postdoctoral predoctoral and postdoctoral researchers is a valuable use of time? Why or why not?

1. If yes (or it depends), how much time is optimal?
2. If no, what data would convince you that it was a good use of time?
3. What type of programming do you think would be (most) useful?
iii. Are you aware of examples where career and professional development is working well? (here or elsewhere)

1. If yes, what is working at institutions that have successfully launched and maintained career development offices/programs?
iv. Do you have other thoughts to add?
v. Thank you for your time!
vi. We may use themes of this to present, no identifying information will be included, do we have your permission to include themes we discussed here today?

#### External-facing staff (Stakeholder 3)

a. Business Development
b. Alumni Relations
c. Industry Engagement
d. Tech Transfer

##### Sequence of conversation & questions

i. Introduction: reminder of purpose
ii. Which external groups do you typically interact with in your role?
iii. What is the intention/purpose of your majority interaction with external stakeholders in your role?
iv. What are the interest areas of the external stakeholders with whom you primarily interact with?
v. Do the people you interact with have an interest in STEM predoctoral and postdoctoral researchers?
vi. Do you have other thoughts to add?
vii. Thank you for your time!
viii. We may use themes of this to present, no identifying information will be included, do we have your permission to include themes we discussed here today?

#### External stakeholders (Stakeholders 4 and 5)

1. Societies
2. Funding agencies
3. Employers (all types, ex: business & administration, communications, industry research, law, policy & outreach, government)

##### Sequence of conversation & questions

i. Introduction: reminder of purpose
ii. What types of engagement are already in place?
iii. What would make you feel engaged as a partner with an academic institution? Are there modes of interaction that have not been explored or could operate better?
iv. What resources to you believe you have to offer to academic predoctoral and postdoctoral researchers and faculty? Are there any that we’ve been missing?
v. How can our predoctoral and postdoctoral researchers better prepare for entry into your industry?
vi. What would make you want to visit campus to interact directly with our population?
vii. vii) Do you follow the national conversation about career development and outcomes of PhD-trained scientists?
viii. Thank you for your time!
ix. We may use themes of this to present, no identifying information will be included, do we have your permission to include themes we discussed here today?

## Appendix B: Sample Wording for Invitation to participate

### Sample Wording for Detailed Invitation

*NOTE: any italicized content should be tailored to your project*

*(insert university logo/header here or use templated stationary)*

Dear *[INSERT NAME]*,

I hope you are well! I reaching out because I am a member of an NIH Broadening Experience in Scientific Training (BEST) Mastermind Group on Culture Change in Academia around graduate student professional development. I was wondering if I could schedule a time to meet with you to interview you about internal views on *[INSERT STAKEHOLDER SPECIFIC TEXT; e.g., professional development engagement with external stakeholders at our university].* I have a few questions about how you, in your role engage with *[STAKEHOLDER SPECIFIC TEXT; external stakeholders]*, that should take about 15-20 minutes. Thank you so much for considering the request!

[*FOR PERSONAL CONTACTS* - Insert personal statement here and/or generic – e.g., I’m part of a working group with people in my role across the country, we are hoping to speak with external facing stakeholders (such as yourself) to see what the need is for companies when it comes to career exploration. I’ve included more information below; would you be interested in speaking with me more about your experience in your current role?]

*[CONTEXT OF PROJECT & POTENTIAL OUTCOMES]* In order to provide resources to predoctoral and postdoctoral researchers, we first need to determine whether our current understanding of ‘the value of a PhD’ is accurate, from the point of view of multiple stakeholder groups; perhaps we are missing important aspects/beliefs that have not been adequately appreciated or explored. We will approach different stakeholders and pose a specific set of questions that explore their relationship and vision of interacting with industry and academia and, specifically, with predoctoral and postdoctoral researchers including how academic institutions and staff can better support those interactions. What we learn from them will help us to better support and develop relevant resources for our predoctoral and postdoctoral researchers.

Example questions include: *[INSERT STAKEHOLDER-SPECIFIC QUESTIONS HERE]*

- Do the people you interact with have an interest in STEM predoctoral and postdoctoral researchers?
- Which external groups do you typically interact with in your role?
- What is the intention/purpose of your majority interaction with external stakeholders in your role?
- What are the interest areas of the external stakeholders with whom you primarily interact with?

Best regards,

*[INSERT INTERVIEWER NAME and CONTACT INFO]*

### Sample Wording for Casual Invitation

*NOTE: any italicized content should be tailored to your project*

*(insert university logo/header here or use templated stationary)*

Dear *YYY*,

I am [*a member of an NIH Broadening Experiences in Scientific Training Mastermind Group working on graduate student professional development*]. I am reaching out about

[*e.g. a research study our group is conducting*] in which I hope you will consider participating.

The purpose of this study is [*to determine whether our current understanding of ‘the value of a PhD’ is accurate from the point of view of multiple stakeholder groups, (e.g. predoctoral and postdoctoral researchers, faculty, academic administration, career services offices, tech transfer offices)*]. The group involved in this study are *graduate education professionals across multiple universities in the country*]. We will [*conduct interviews with specific sets of questions that explore your opinion, relationship, and vision of interactions between industry and academia*]. What we learn will help us to [*better support development of our predoctoral and postdoctoral researchers for what the workforce needs*]. Interviews will last approximately [*30 minutes*]. [*Our interview will ask about aspects of career and professional development (e.g. what types of engagement are already in place, how can our predoctoral and postdoctoral researchers better prepare for entry to your industry, what resources you may have to offer)*]. Your feedback is extremely valuable to us, and I would greatly appreciate it if you would be willing to participate. Please let me know if this is of interest to you.

Thank you,

*Your name*

*Contact information*

## Appendix C Results with fewer than 5 mentions

### Stakeholder 1 – Internal Pre- and Postdoctoral Researchers

#### Prestige- recruitment (4 mentions)

Predoctoral and postdoctoral researchers feel that having access to offerings can help with their recruitment to companies. Gael succinctly communicated that professional development programs help build a self-fulfilling good reputation for the institution that interests companies, and in turn improves predoctoral and postdoctoral researcher’ track records of landing jobs. Academic-career focused Gunnar also noted that these programs may have unforeseen benefits for pre- and postdoctoral researchers such as facilitating collaboration with industry and promoting translational research.

#### Growth/challenges (4 mentions)

Pre- and postdoctoral researchers acknowledged that career exploration can be challenging but leads to growth. This theme is captured in Glenn’s statement that researchers have to try to push boundaries. Predoctoral and postdoctoral researchers recognize there are many unknowns with regards to CPD and how it is better to go into the process aware of how hard it will be. Pre- and postdoctoral researchers offered several examples of programming that they think would be most useful, ranging from a condensed and intensive multi-day workshop to online training such as sessions the NIH Office of Intramural Training and Education (OITE) offers. Another suggestion, repeated in multiple contexts, was that biomedical science programs model professional schools such as Business and Applied Science programs. Some felt that one-on-one coaching was the most efficient approach and a number mentioned specific skills that they felt would be the most useful including people management, budgeting, negotiations, and personal branding.

#### Faculty permission (4 mentions)

Pre- and postdoctoral researchers communicated their concern about faculty buy-in, and, as Glenn suggested, the need to introduce the concept of professional development early to faculty so they are more comfortable with career exploration. Gunnar, a non-user, suggested that faculty approval to participate should not be required. Of note was the sentiment that even help with academic skills such as grant writing assistance from one’s mentor should be provided during a structured time, apart from time for other professional development activities.

### Stakeholder 2 – Internal Faculty and Administrators

#### Cycle of positive fulfilment for program (3 mentions)

Individuals commented that future applications to the university will benefit from the existence of a good professional development program, since many prospective students look for these opportunities. Fabio captured this sentiment, commenting that a professional development program is valuable for faculty to attract good people to the lab.

### Stakeholder 3 – Internal Partners-External Facing Staff

#### Benefit to predoctoral and postdoctoral researchers (4 mentions)

In these external-facing offices’ views, the primary reason to engage with external stakeholders is to provide exposure. As Simha shared, she wants to expose researchers to perspectives other than her own, and the diversity of perspectives and options available on campus are enhanced by industry interactions.

#### Pay it forward (3 mentions)

Some external-facing staff find that external partners are interested in helping in any way they can, in an effort to give back to institutions, while others find industry professionals are interested in helping, in order to be part of an exciting new professional development activity. Shanice mentioned that since external stakeholders understand what students need to be successful, they believe can help the next generation, and in fact wish that they had these types of programs when they were pre- and postdoctoral researchers.

Although not mentioned frequently enough to constitute a theme, a few internal-external partners mentioned that *prestige (2 mentions)* played a role in their engagement with academia. Some external partners are interested in developing relationships to create legacies in their names at the institutions. Saachi explained that external stakeholders feel they gain credibility by interacting with the university and being considered an asset.

#### Support student needs (4 mentions)

The general opinion is that external stakeholders have an interest in STEM pre- and postdoctoral researchers because they understand and support faculty and pre- and postdoctoral researcher needs. Shanice commented that they want to contribute to the pre- and postdoctoral researcher professional development and academic experience. They recall their own educational experience and reflect upon what is needed.

#### Communication coordinator (4 mentions)

Several external-facing staff thought that there should be a coordinated effort to engage with external partners. In particular, Shanice thought that their office of alumni engagement should be used to connect with alumni, and coordinate communication.

#### Alumni engagement role (3 mentions)

Many external-facing office initiatives engage alumni, with a fundraising end-goal for institutions. On the other hand, alumni are interested in the prestige or ability to create a legacy, as well as to pay it forward to the next generation of pre- and postdoctoral researchers. Shanice shared that what appears important to each individual alumna/alumnus for engagement is being informed of the expected time commitment and financial request, and expect in return to have networking opportunities and to be connected to the right pre- and postdoctoral researcher for mentoring and/or hiring.

### Stakeholder 4 – External Partners –Non-Profits/ Societies

#### Online resources, guides (4 mentions)

Societies, foundations and non-profit external stakeholders compile several online resources for wider use. A goal of these stakeholders is to create and share among the community, and as Ellis stated, they hope more universities will engage with the resources they have compiled.

#### Experiential learning (4 mentions)

A resource many institutions are looking for is experiential learning opportunities, and some societies, foundations, and non-profits offer this resource. Models include short-term fellowships, or longer period training mechanisms. Eric described a model within their organization as a mechanism to train leaders. Non-profits advise that pre- and postdoctoral researchers have a “growth mindset” and should use time in graduate school to test the waters to become a leader. A topic that arose only infrequently was *self-exploration (2 mentions)*. An important aspect of career planning and exploration is the notion of understanding one’s strengths and skills. Eric recommended that pre- and postdoctoral researchers make use of personality assessments, to build awareness of the skills they have that industry seeks.

#### Funding (4 mentions)

Societies, foundations, and non-profits face funding limitations. Though they are keen to encourage careers at their organization, they may have limited resources for fellowships or research. Ellis described how the rate of funding they receive annually has been stagnant, and hence, unfortunately, fewer awards are given out each year. Importantly, these partners encourage that the affiliation between non-profits and academia should focus on relationships and resources, other than financial reliance.

#### Connect with target audience (3 mentions)

External entities struggle with knowing whom to contact at institutions to ensure they are reaching the appropriate audience. Ebony recommended that universities have a one-stop shop, so external entities know whom to contact.

#### Flexible, creative models (3 mentions)

Another challenge faced by these stakeholders is interacting within the strict design of educational programs at universities. They recommend flexibility to allow more exploration time for students; Ellis highlighted the need to propose different ways of training to prepare students for their futures, and to create and track these models with evidence-based outcomes and effectiveness research. Non-prescriptive models for graduate learning could open doors for more engagement with non-profits and foundations. Relatedly, Evan communicated the desire to talk to faculty about the importance of career exploration.

#### Listen with humanity, broaden diversity (4 mentions)

A theme emerged from multiple conversations that being an inclusive leader is crucial, and having that outlook is important for pre- and postdoctoral researchers. Eric mentioned the need to listen to the majority and also historically under-represented minority groups, while Emily discussed the need to learn how to support underrepresented groups to help the next generation of scientists. Ellen highlighted the need to pause and learn about others, and to not always center one’s conversation around an individual’s point of view.

Although mentioned infrequently among these participants, and hence not classified as a theme of its own, a key concept to finding the right career path is *networking and engagement opportunities (2 mentions)*; Ebony advised pre- and postdoctoral researchers find ways to connect beyond conferences such as SACNAS, and suggested the need for regional conferences centered around graduate careers to help expose pre- and postdoctoral researchers to opportunities.

Some pieces of advice were provided to researchers, such as *get involved in professional societies (1 mention)* to expand their spheres and take on leadership roles. Moreover, *defining success (1 mention)* of the relationship should be left up to the fellows or pre- and postdoctoral researchers themselves since they know what metrics are important to them. As Ebony advised, fellows should learn to identify their own success measures.

> “Definition of success is defined by the individual; the PhD is a key to the door but what door do you want to open?”

A few individuals mentioned that they did not want to come to campus but rather wanted to *take the predoctoral and postdoctoral researchers off campus (2 mentions)* to experience advocacy in the government or how their non-profit operates in the real world.

*A few varied challenges* were identified infrequently by external partner organization stakeholders but should be taken into consideration. One challenge identified was that an organization’s (e.g. non-profit) *offerings should be integrated* into training *(2 mentions).* Although flexibility is necessary as mentioned above, some participants were keen, and even preferred to embed their resources in the graduate curriculum, to ensure all graduate student pre- and postdoctoral researchers have access.

External partners wish to *engage at all levels with academic institutions (2 mentions).* Strategic planning requires engagement by all members of an academic institution; hence societies, foundations, and non-profits must prioritize the amount of engagement necessary at each level, to achieve their goals, and institutions must be willing to engage.

An external partner mentioned the challenge that the *wrong people are making decisions (1 mention)* at some of these external organizations, and it appeared that once PhD-level employees were involved in the decision-making process, the offerings they provided to academia were much more relevant.

Finally, another challenge faced by external partners is the need for more industry mentors. Ebony describes the difficulty in getting enough mentors for the fellows program at their organization.

### Stakeholder 5 – External Employers -Industry

#### Bidirectional partnerships (4 mentions)

Some employers even rely on universities to facilitate their research and development. As Ira pointed out, pre- and postdoctoral researchers generate early discovery and pre-clinical data. Companies see this as a two-way relationship that benefits both parties.

#### Advisory and feedback roles (3 mentions)

Relationships between academic institutions and companies create and strengthen connections between the two. For example, Ira suggested industry can conduct focus groups to learn how to better tailor offerings such as internships to students, and while Ivy commented that industry can, and sometimes does, share with faculty how skills transfer to industry.

#### Learn industry-relevant skills (4 mentions)

External stakeholders shared an important opinion: they do not think that the graduate school curriculum prepares pre- and postdoctoral researchers sufficiently for industry since the goals of research are different in the two sectors. Ira emphasized this idea, saying academics must learn how to better design experiments of relevance to industry. Since the goal of biotech is to commercialize rather than publish, experimental criteria are very different.

#### Recalibrate importance (3 mentions)

Industry and academia place weight on different metrics and therefore predoctoral and postdoctoral researchers need to shift their focus when applying for jobs. Iris bluntly stated that not all individuals in industry value publications when vetting new hires. Instead predoctoral and postdoctoral researchers should focus on how the transferable skills they have can help the company move a product through the pipeline which is what is important to industry.

#### Grants and academic collaborations (3 mentions)

Irina keenly noted that industry is to be viewed as a collaborator, and not a piggy bank. Industry employs top scientific minds, who are eager to collaborate with academia. Their goal, however, is not to help solve problems in research, but rather to advance their product to commercialization.

#### Point of contact (3 mentions)

Ian pointed out that the relationship between industry and academia needs longevity and suggested that universities should identify an individual program director who assigns or recruits students. The complexity of the organizations often creates difficulties for either party (companies and universities) to know who the appropriate point person is in the other party. Relatedly, to maintain longevity, Ian also impressed upon the importance of tracking the impact of an interaction or event.

#### Faculty culture change (4 mentions)

An oft-thought opinion was voiced by Ivan when he commented that students shouldn’t feel like they are operating behind their advisor’s back. To aid in faculty culture change, Ivy advised that CPD practitioners share with faculty how skills transfer to industry and offer perspectives. Irina, aware of management hierarchies and dynamics, advised making changes from the top, by approaching department chairs or deans to be open-minded to industry career outcomes for their graduates. Faculty culture needs to change to accept different career paths.

#### High-impact events (3 mentions)

Iris described how more comprehensive engagement with faculty during their visits (e.g., larger events can involve faculty beyond the event, networking opportunities, informal interactions, one-on-one meetings) helps make industry partners feel welcome. Industry has interest if a university hosts events that guarantee large pre- and postdoctoral researcher attendance, so they have maximal impact for their visit.

#### Match-ups (3 mentions)

Ian mentioned the value of listservs or student groups who are interested in various companies’ work, as these lists can be shared with appropriate companies as a recruiting tool. Companies would be interested in interacting if they are assured to find recruits who match up as the appropriate kind of candidate.

#### Open-minded to industry (3 mentions)

While culture change is important, issues pertaining to intellectual property constraints, or access to departments involved in translational research were described as a challenge.

*A few other comments* represented important opinions, but were not repeated across interviews, and are hence not classified as themes. A participant described they had no desire to visit campus, but instead preferred to host site visits for pre- and postdoctoral researchers. Another described limited bandwidth at their organization to devote time and resources to university relations.

